# Semaphorin signaling restricts neuronal regeneration in *C. elegans*

**DOI:** 10.1101/2021.11.13.468479

**Authors:** MB Harreguy, Z Tanvir, E Shah, B Simprevil, TS Tran, G Haspel

## Abstract

Extracellular signaling proteins serve as neuronal growth cone guidance molecules during development and are well positioned to be involved in neuronal regeneration and recovery from injury. Semaphorins and their receptors, the plexins, are a family of conserved proteins involved in development that, in the nervous system, are axonal guidance cues mediating axon pathfinding and synapse formation. The *Caenorhabditis elegans* genome encodes for three semaphorins and two plexin receptors: the transmembrane semaphorins, SMP-1 and SMP-2, signal through their receptor, PLX-1, while the secreted semaphorin, MAB-20, signals through PLX-2. Here, we evaluate the locomotion behavior of knockout animals missing each of the semaphorins and plexins and the neuronal morphology of plexin knockout animals; we described the cellular expression pattern of the promoters of all plexins in the nervous system of *C. elegans;* and we evaluated their effect on the regrowth and reconnection of motoneuron neurites and the recovery of locomotion behavior following precise laser microsurgery. Regrowth and reconnection were more prevalent in the absence of each plexin, while recovery of locomotion surpassed regeneration in all genotypes.

## Introduction

During neurodevelopment, growth factors and guidance cues regulate dendrite morphogenesis, axon growth cone initiation and navigation, axon elongation and target recognition, but their effects are less pronounced in the adult nervous system. Studying their role in the context of adult regeneration and recovery could provide insight into the molecular and cellular response to injury (Chen et al., 2011; Chisholm et al., 2016).

The semaphorins are a family of glycosylated proteins that were first characterized for their role in the development of the insect and avian nervous systems as axonal guidance cues but were later found in a variety of other tissues and organisms (Alto and Terman, 2017; Junqueira Alves et al., 2019). All semaphorins have a distinctive 500 residue long N-terminal domain, known as the Sema domain. This domain, which is a seven-blade beta-propeller, with each blade formed by four anti-parallel beta-strands (Gherardi et al., 2004), is exclusive to semaphorins and their receptors, the plexins, where it mediates semaphorin dimerization and receptor binding. Eight classes of semaphorins are phylogenetically conserved in nematodes, flies, chick, mammals, and viruses, with three classes of smaller proteins that are secreted and five classes that are membrane-bound by a transmembrane domain or a glycosylphosphatidylinositol (GPI) link (Alto and Terman, 2017; Junqueira Alves et al., 2019). Correspondingly, four classes of plexins are conserved in invertebrates and vertebrates (Tamagnone et al., 1999; Negishi et al., 2005). All plexins are transmembrane proteins with an extracellular Sema domain that mediates semaphorin binding and signaling, either by themselves or with a neuropilin co-receptor, in the case of the secreted class 3 semaphorins in vertebrate (Negishi et al., 2005; Pascoe et al., 2015).

In mammals, semaphorins and their receptors, neuropilins and plexins, were originally described as guidance cues for neuronal growth cones aiding axons to their targets by acting as chemorepellents (Kolodkin and Tessier-Lavigne, 2011). More recently, semaphorins have been implicated in multiple key roles of neural circuit assembly during neurodevelopment (Yoshida, 2012; Koropouli and Kolodkin, 2014). For example, the mammalian secreted semaphorin, SEMA3A, is involved in various neurodevelopmental processes in the mouse, including repelling dorsal root ganglion sensory axons, promoting basal dendrite elaboration in cortical pyramidal neurons, and pruning of hippocampal axons (Bagri et al., 2003; Yaron et al., 2005; Mlechkovich et al., 2014; Danelon et al., 2020). Another well studied secreted semaphorin, SEMA3F, and its receptor Neuropilin-2, are also involved in axon guidance, synaptic plasticity, and refinement, as well as in restraining the excess of dendritic spines on apical dendrites of cortical neurons and regulating inhibitory interneuron numbers in the hippocampus (Tran et al., 2009; Riccomagno et al., 2012; Riccomagno and Kolodkin, 2015; Assous et al., 2019; Eisenberg et al., 2021). As the mediators of semaphorin signaling, the plexins are involved in axon guidance, synapse and dendrite formation, axonal pruning and synaptic stability (Shen and Cowan, 2010; Limoni, 2021).

In accordance with their role in neurodevelopment, semaphorins could be involved in axonal regeneration after injury (Fard and Tamagnone, 2021). For example, SEMA3A expression levels increase after injury in the spinal cord and cerebral cortex (de Winter et al., 2002; Hashimoto et al., 2004) and regenerating axons avoid areas with high SEMA3A expression (Pasterkamp and Verhaagen, 2001). Accordingly, a SEMA3A-specific inhibitor improved axon regeneration and spontaneous hind leg movement after spinal cord transection (Kaneko et al., 2006). Plexin expression and function in response to injury varies depending on the type. Plexin A family members increase their expression after axonal injury in facial motoneurons and rubrospinal neurons contributing to the role of semaphorins in restricting regeneration (Spinelli et al., 2007). On the other hand, PlexinB2 is upregulated after spinal cord injury in glial cells proximal to the injury site and is required for wound healing and recovery (Zhou et al., 2020).

The *Caenorhabditis elegans* genome encodes for only three semaphorin and two plexin homologues. Of those, PLX-1 binds the two membrane-bound semaphorins (SMP-1 and SMP-2), while PLX-2 binds the only secreted semaphorin (MAB-20; Fig. 1A; (Ginzburg et al., 2002; Nakao et al., 2007). Both membrane-bound and secreted semaphorin-plexin systems are involved in development; semaphorins guide ventral enclosure (Ikegami et al., 2012), and regulate epidermal morphogenesis (Ginzburg et al., 2002; Ikegami et al., 2012) as well as vulva and tail-rays morphogenesis in the hermaphrodite and males, respectively (Dalpe et al., 2012). In the nervous system, membrane-bound semaphorin signaling (the *plx-1/smp-1/smp-2* pathway) is necessary for synaptic tiling in two DA motoneurons in the tail (Mizumoto and Shen, 2013) and for guidance of the long axons of mechanosensory neurons (Ginzburg et al., 2002). Secreted semaphorin signaling (via the *plx-2/mab-20* pathway) contributes to motoneuronal axon guidance; eliminating this pathway, when not embryonic lethal, causes defasciculation of the ventral nerve cord (VNC; 17% of surviving *mab-20* knockout animals) and axon misguidance in DA and DB motoneuron classes (4% of surviving *mab-20* knockout animals;(Roy et al., 2000).

**Figure 1.**
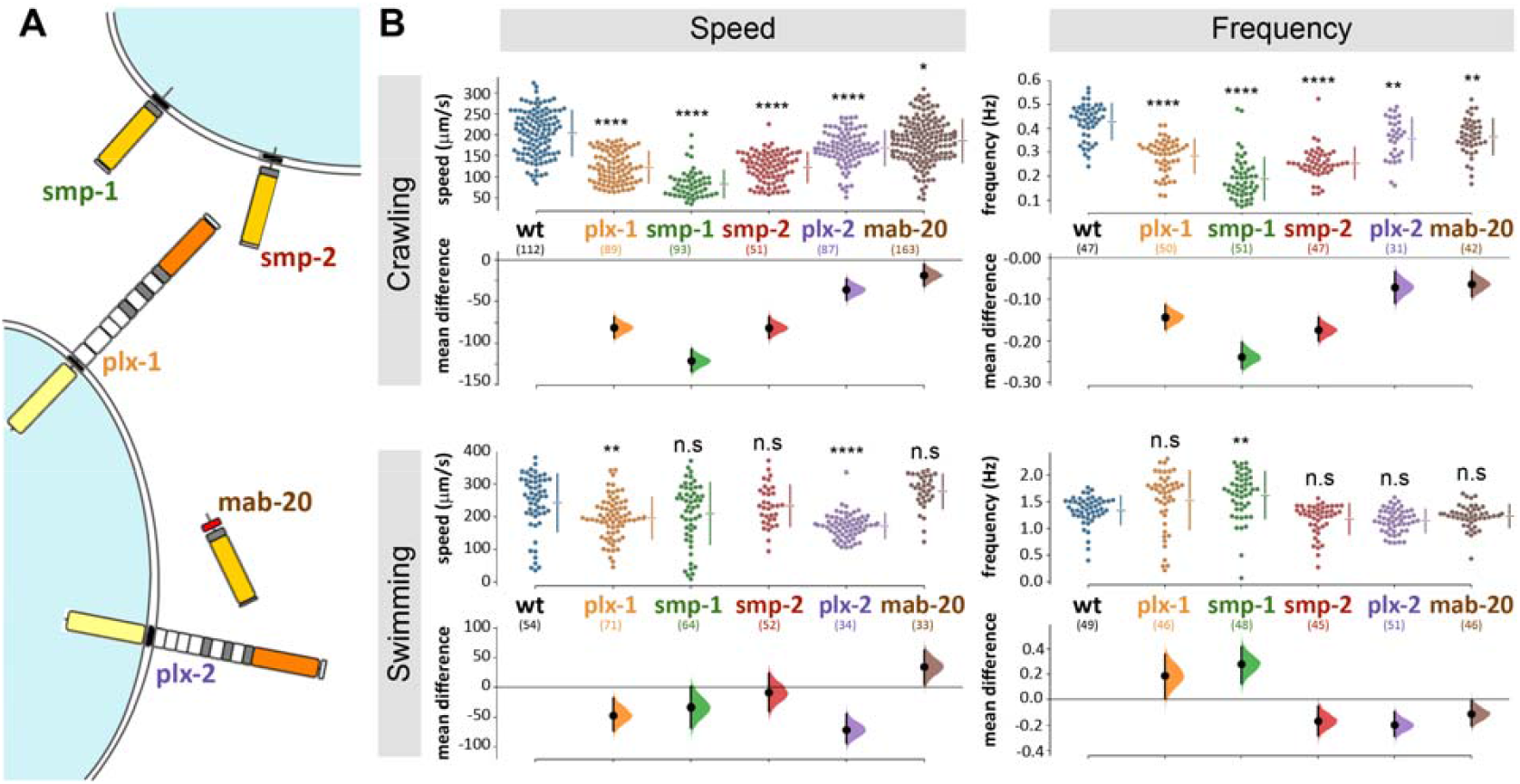
*C. elegans* semaphorin system comprises only three ligands and two receptors and omitting any one component affects locomotion. A) Semaphorin signaling system of *C. elegans*. The membrane bound semaphorins *smp-1* and *smp-2* signal through *plx-1*, while the secreted *mab-20* signals through *plx-2* (molecular diagrams adapted from Junqueira Alves et al., 2019). B) Mutant strains with knocked out semaphorins or plexins are significantly different from wild type when crawling (locomoting on agar) or swimming (locomoting in liquid media). The largest difference was in *smp-1*(ko) animals while *mab-20*(ko) animals are only different for lower undulatory frequency during crawling. Data points are mean absolute translocation speed or frequency to both direction of locomotion of analyzed trajectories; n.s. p>0.05, ^*^p<0.05, ^**^p<0.01, ^***^p<0.001, ^****^p<0.0001; one-way ANOVA with Tukey’s multiple comparisons test post hoc; in parentheses are the number of analyzed trajectories from 20-25 animals for each genotype.

*C. elegans* is a well-established model for neuronal regeneration and most of its neurons are able to regenerate after precise laser microsurgery (60%) and in some cases reestablish functional connections (Yanik et al., 2004; Ghosh-Roy and Chisholm, 2010; Neumann et al., 2011; Harreguy et al., 2020; Harreguy et al., 2022). **Here we take advantage of the small number of plexins in *C. elegans* and the capability to precisely disconnect single neurites in intact animals, to investigate the role of semaphorin signaling in neuroregeneration *in vivo*. We describe the neuronal expression of the plexin receptors and the effect of their absence on neuronal regeneration and recovery of locomotion behavior**.

## Methods

### Strains and transgenics

We maintained *C. elegans* strains under standard laboratory conditions on nematode growth medium (NGM: 0.25% Tryptone, 0.3% Sodium Chloride, 1 mM Calcium Chloride, 1 mM Magnesium Sulfate, 25 mM Potassium Phosphate (pH 6.0), 5 μg/mL Cholesterol, 1.5% Aga) agar plates with OP-50-1 *Escherichia coli* bacterial lawn at 15°C (Stiernagle, 2006), without antibiotics. All animals used in the experiments were hermaphrodites.

We acquired semaphorin and plexin mutants from Caenorhabditis Genetics Center (CGC) or the *C. elegans* National Bioresource Project of Japan (NBRP): ev778 (*mab-20*, null), tm729 (*plx-2*, null), ev715 (*smp-1*, null), ev709 (*smp-2*, null), tm10697 (*plx-1*, null), and evIs111 ([F25B3.3::GFP + dpy-20(+)], pan-neural GFP expression). To allow imaging and microsurgery, we crossed males of NW1229 (evIs111), induced by 10-minute exposure of L4 larvae to 10% ethanol (Lyons and Hecht, 1997), with null-mutant hermaphrodites to obtain knockout animals expressing GFP in the entire nervous system: TOL55 (ev715, evIs111, outcrossed x6), TOL57 (ev709, evIs111, outcrossed x6), TOL59 (tm10697, evIs111, outcrossed x1), and TOL62 (tm729, evIs111, outcrossed x1). All strains were verified by PCR upon arrival, after crosses, and at the end of the study. All generated strains and primer sequences for genotyping will be deposited with the CGC.

The reporter strain for *plx-1p*::EGFP (NW2339, 2,621 bp sequence immediately 5′ to the ATG start codon cloned into the multiple cloning site of pPD95_77; Dalpé et al., 2004) and *plx-2p*::GFP (NW1693, 4,529 bp sequence immediately 5′ to the ATG start codon cloned into the multiple cloning site of pPD95.75) were generous gifts from Dr Joseph Culotti (University of Toronto, Mt Sinai Hospital) and Dr Richard Ikegami (UC Berkeley), respectively. For unambiguous identification, we crossed each reporter strain with a NeuroPAL transgenic strain (OH15495; Yemini et.al 2021).

### Locomotion analysis

We tracked locomotion behavior of multiple animals over an agar surface (1.7% in NGM buffer), without food, as well as in liquid (NGM buffer). We recorded videos with a static multi-worm tracker, composed of three major parts, from top to bottom: (1) a CMOS camera (acA4024-29um, Basler) mounted with a fixed focal length lens (C Series 5MP 35 mm 2/3”, Edmund Optics), and an infrared cut-off filter (SCOTT-KG3 M25.5×0.5, Edmund Optics); (2) a specimen stage for plates or slides; (3) a collimated Infrared LED light source (M850L3 and COP1-B, Thorlabs).

One day before the experiment, we transferred animals of the fourth larval stage (L4) onto a new plate with healthy OP-50-1 bacterial lawn. Ten to fifteens minutes before tracking, animals were transferred onto a 30 mm agar plate with no food or a 150 μL drop of NGM buffer, placed on a microscope slide. During tracking, animals moved freely, and we recorded multiple 25 Hz 15-second videos using Pylon Viewer (Pylon Camera Software Suite, Basler). We analyzed the videos with Tierpsy worm-tracker (Javer et al., 2018) that can track multiple animals and extract up to 726 features for each tracked trajectory. We used the Tierpsy post-processing user interface to merge tracked sections (trajectories) if those were erroneously split by the automatic tracking, and we rejected any trajectory shorter than 3 s, as well as ambiguous cases of animal proximity. Recording and Tierpsy analysis were done by undergraduate researchers, blinded to the animals’ genotype and injury condition. We analyzed the HDF5 output file produced by Tierpsy with a MATLAB script (code available upon request) to collect the mean speed and frequency values for each trajectory and then plotted the data and estimated confidence intervals between each group and its control with a freely available software for Estimation Statistics (https://www.estimationstats.com; (Ho et al., 2019) that focuses on the magnitude of the effect (the effect size) and its precision. We also present statistical significance calculated with a two-sided permutation t-testto compare sham vs injured groups, or one-way ANOVA with Tukey’s multiple comparisons test post hoc to compare genotypes (GraphPad Prism v9.2), included as p values in the text and as asterisks that denote levels of significance. We routinely use this tracking system to evaluate and compare wild type, injured, and uncoordinated mutant strains. We tracked all the knockout, transgenic, and wild type strains without injury to assess their baseline locomotion parameters. Further, we tracked locomotion to assess recovery 6, 12, and 24 hours after microsurgery. For comparison, we also quantified locomotion parameters of sham-surgery groups for each genotype and time point. We treated the sham-surgery groups through the same protocol (including cooling and immobilization, see below), except for the exposure to the laser beam.

### Expression and neuronal morphology analysis

To reduce autofluorescence and straighten the animals we incubated fourth stage larvae (L4) in M9 buffer for 90 m and washed in the same buffer three times, incubated in 1 mM Levamisole (a paralytic nicotinic agonist, Sigma Aldrich) for 15 m, and fixed overnight at 4°C in 10% formalin solution, neutral buffered (SIGMA), then washed and mounted with Fluoromount-G (EMS), and allowed the slides to dry for at least 24 h before imaging. We used a laser scanning confocal microscope (Leica SP8; microscope: DM6000CS; objectives: Leica 40x/NA1.30 HC PL APO oil or Leica 63x/NA1.40 HC PL APO oil, with lateral resolutions of 223 nm and 207 nm respectively; laser lines: 405 nm, 561 nm, and 488 nm). We collected multiple optical slices (thickness optimized by the confocal software, ranging 0.343 - 0.345 μm for the 63X objective, and 0.410 - 0.422 μm for the 40X objective). To analyze morphology and cellular expression we constructed the maximum intensity projections for at least 10 animals of each strain and, in some cases, processed images to reduce background noise via the Leica Application Suite (LASX) software.

For unambiguous identification of VNC motoneuronal expression, we crossed each transcriptional reporter strain with a NeuroPAL transgenic strain and imaged the F1 progeny that express both transgenes. The NeuroPAL strains express an invariant color map across individuals, where every neuron is uniquely identified by its color and position (Yemini et.al 2021). We identified 29 motoneurons in 3 animals and rejected 3 motoneurons that expressed GFP but their location and NeuroPAL colors were ambiguous.

### Laser Microsurgery

For laser microsurgery and associated microscopy, we mounted *C. elegans* hermaphrodites at L4 stage by placing them in a drop of ice cold, liquid 36% Pluronic F-127 with 1□mM levamisole solution and pressed them between two #1 coverslips (Melentijevic et al., 2017). We brought the coverslips to room temperature, to solidify the Pluronic F-127 gel and immobilize the animals. We used a Yb-fiber laser (100 pulses at 10 kHz repetition rate) to cut a single neurite with submicron precision and no discernable collateral damage (Harreguy et al., 2020; Harreguy et al., 2022). We took images immediately before and after the lesion to visually verify the microsurgery. In some cases, multiple laser exposures were necessary to disconnect a neurite. We disconnected the ventral-dorsal commissures of all motoneurons that we were able to identify by their relative position (at least 6 per animal), at about 45 μm away from the VNC. We assessed neuronal regeneration 24□hours (following most regeneration studies in *C. elegans*, since Yanik et al., 2004) after microsurgery on the same microscope and imaging system in at least 6 neurons per animal in at least 15 animals for each condition. We considered neurites regrown when a new branch was observed extending from the proximal segment of the injury site (Harreguy et al., 2020; Harreguy et al., 2022). When the branch extended to the distal segment or the target of the pre-injury neurite, we considered it regrown and reconnected. We used Fisher Exact on 2×3 contingency table to compare the fraction of observed neurites that regrew or reconnected. We used ImageJ (FIJI v.1.52) and LASX (Leica) for image processing and visualization, and Prism (GraphPad v.9.2.0) for statistical analysis and plotting.

## Results

### C. elegans animals that do not express functional semaphorins or plexins exhibited altered locomotion patterns

We analyzed the contribution to locomotor behavior of each of *C. elegans* three semaphorins and two plexins (Fig. 1A) by comparing the speed and frequency of locomotion of knockout (ko) mutant strains to that of wild type animals. During crawling on agar (Fig. 1B), all strains translocated significantly slower compared to 204±54 μm/s of wild type (speed and p values were: *plx-1* 123±37, p<0.0001; *smp-1* 83±33, p<0.0001; *smp-2* 123±35, p<0.0001; *plx-2* 168±41, p=0.0011; *mab-20* 186±51, p=0.0016); and the undulation frequency of all strains was reduced compared to 0.43±0.08 Hz of wild type (frequency and p values were: *plx-1* 0.29±0.07, p=0.0497; *smp-1* 0.19±0.09, p<0.0001; *smp-2* 0.25±0.06, p<0.0001; *plx-2* 0.36±0.09; *mab-20* 0.36±0.08). Relative to crawling, swimming speed and frequency were less affected by the absence of plexins or semaphorins (Fig. 1B), only *plx-1(ko)* and *plx-2(ko)* animals translocated slower than 243±88 μm/s of wild type (speed and p values were: *plx-1* 196±63, p=0.003; *smp-1* 209±94; *smp-2* 234±63; *plx-2* 172±38, p<0.0001; *mab-20* 277±52); only *smp-1(ko)* animals undulated at higher frequency compared to 1.34±0.27 Hz of wild type (frequency and p values were: *plx-1* 1.53±0.55; *smp-1* 1.62±0.44, p=0.0014; *smp-2* 1.17±0.29; *plx-2* 1.14±0.22; *mab-20* 1.23±0.22). The largest reduction of crawling speed and frequency was in *smp-1(ko)* animals that were also the only genotype to exhibit a change (increase) in undulation frequency during swimming.

We focused further analysis on the plexins (*plx-1* and *plx-2*), because as the only receptors, segregating membrane-bound and secreted pathways, they provide a comprehensive and specific manipulation of these pathways, as well as the identity of the cellular targets (Fujii et al., 2002).

### Gross neuronal morphology was unaffected by the absence of PLX-1 and PLX-2

We used confocal microscopy to image at least 5 intact four instar (L4) larvae of each plexin-knockout and wild type strain, expressing pan neuronal green fluorescent protein (GFP), with emphasis on neuron-rich areas around head, tail, the ventral nerve cord, pharynx, and vulva, and particularly at the commissures of motoneurons (Fig. 2). We did not observe any morphological differences between mutant and wild type animals in any of these regions.

**Figure 2.**
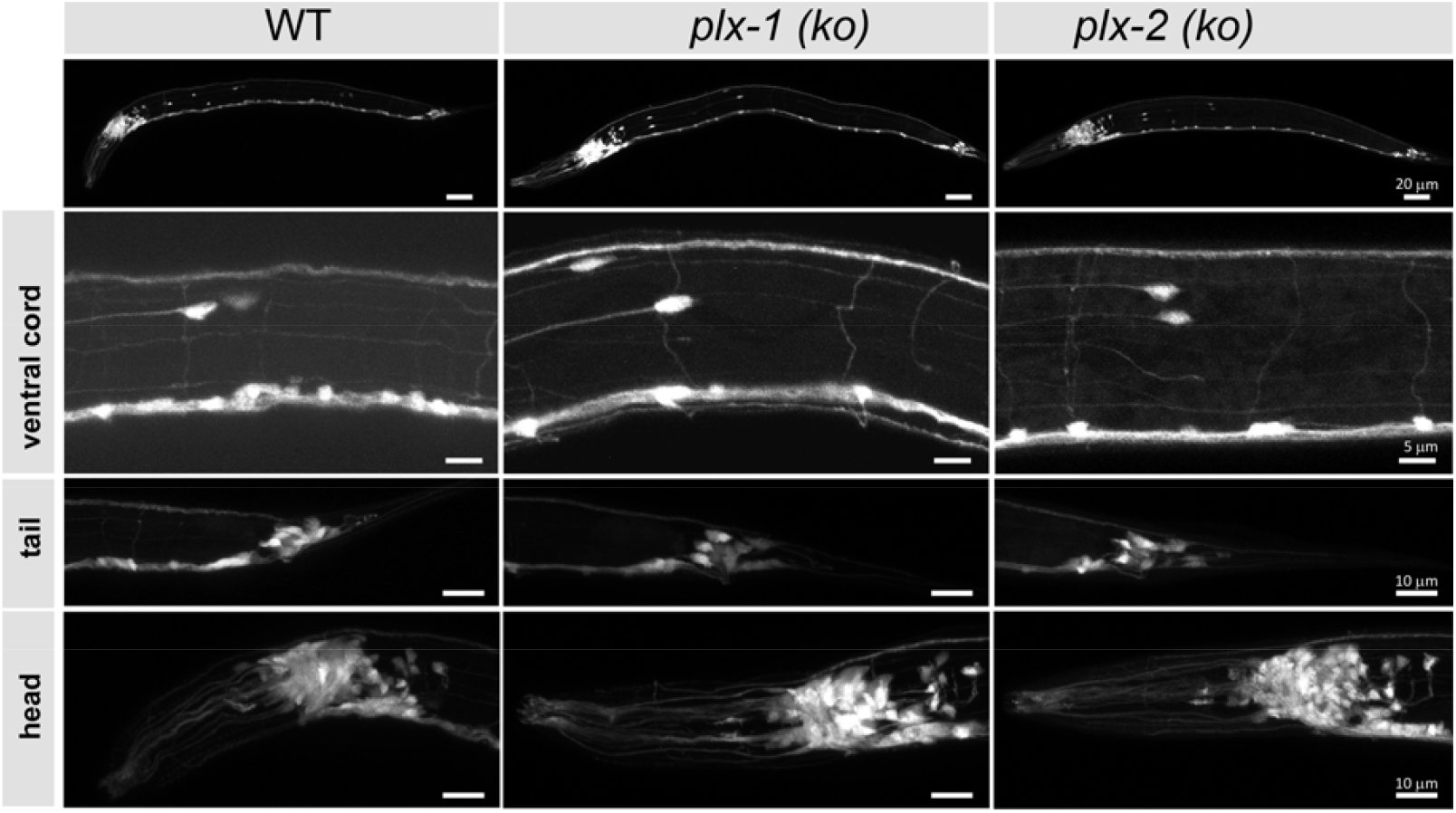
Neuronal morphology of plexin knockout strains is comparable to wild type. The nervous systems are visible via pan-neuronal GFP in neuron-rich areas (VNC, head, and tail ganglia) of wild type (WT) and knockout mutant animals (*plx-1*(ko) and *plx-2*(ko)), as well as the entire animals (top), to look for gross neuromorphological differences. We did not observe differences between wild type and mutant strains. N>5 animals for each strain. Scale bar = 20 μm (whole animals), 5 μm (VNC), and 10 μm (bottom panels).

### Motoneuronal expression of PLX-1 and PLX-2

We imaged transcriptional reporters for *plx-1p* and *plx-2p* in order to identify their neuronal expression in the ventral nerve cord (VNC). GFP under the *plx-1p* promoter (Fig. 3AB) was mostly expressed in non-neuronal tissue including the pharyngeal muscle, the body-wall muscle in the head and along the body, and vulva muscle. We did not find expression in the nervous system of *plx-1*p::GFP, although a translational reporter was reported to express in the axon of a motoneuron at the base of the tail, namely DA9, of the embryo and L1 larva (Mizumoto and Shen, 2013). GFP under the *plx-2*p promoter was expressed by neurons in the head and tail (Fig. 3C), as well as in motoneuron in the VNC (Fig. 3D). Most expressing motoneurons were AS and DA classes (14 and 9, respectively, from 3 animals), 6 motoneurons of other classes, namely DB (3), VA (2), and VB (1) also expressed GFP. Both AS and DA extend commissures that were the targets for microsurgery, from the VNC to the dorsal nerve cord on the opposite side of the animal.

**Figure 3.**
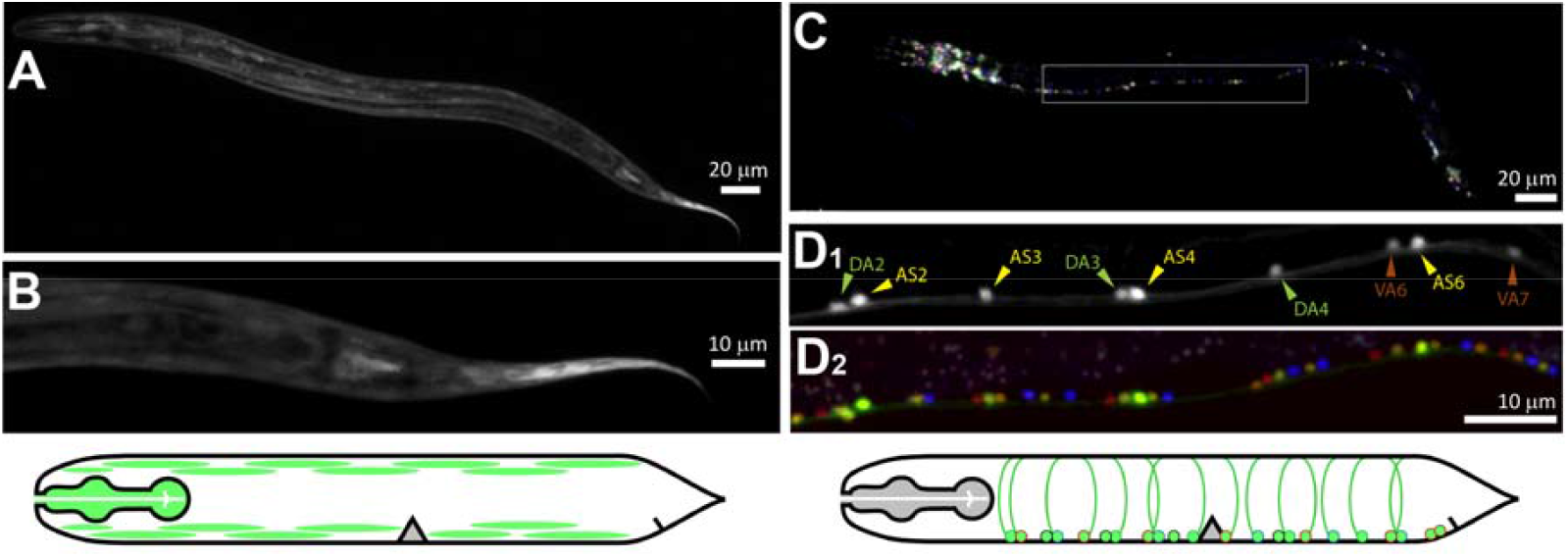
PLX-1 is expressed in non-neuronal tissue, while PLX-2 is expressed in excitatory motoneurons. AB) Green fluorescent protein (GFP) driven by *plx-1p* promoter expressed in non-neuronal tissue such as the pharynx, body-wall muscle. C) GFP driven by *plx-2p* promoter expressed mostly in AS and DA motoneurons and in a few DB, VA, and VB motoneuron. D_1_) Examples of DA2-4, AS2-6, and VA6-7 that were identified with co-expressed NeuroPAL (D_2_). Images in A and B are from different animals. Scale bars are 20 μm (AC) and 10 μm (BD).

### Neurites of plexin knockout mutants regenerate more than wild type after laser microsurgery

We disconnected 156 commissural neurites of motoneurons of wild type and plexin knockout mutant animals with laser microsurgery (Harreguy et al., 2020; Harreguy et al., 2022). These lateral processes extend to connect the ventral and dorsal nerve cords and when multiple processes are disconnected, locomotion is impaired (Yanik et al., 2004). When we examine the same neurite after 24 hours, some regrew by sprouting a growth cone from the proximal segment and some of those reconnected to the distal segment or the dorsal nerve cord (Fig. 4A). In the wild type, 38 of 73 neurites regrew (0.52±0.11) and only 5 of those (0.07±0.058) reconnected (Fig. 4B). The plexin knockout mutants exhibited significantly more regrowth (p=0.049,), 33 of 47 (0.7±0.13) for *plx-1(ko)* and 26 of 36 (0.72±0.15) for *plx-2(ko)*. Reconnection happened significantly more (p<0.0001) in the plexin knockout strains: in *plx-1(ko)*, 13 of the regrown neurites (0.28±0.13) and in *plx-2(ko)*, 20 of the regrown neurites reconnected (0.56±0.16).

**Figure 4.**
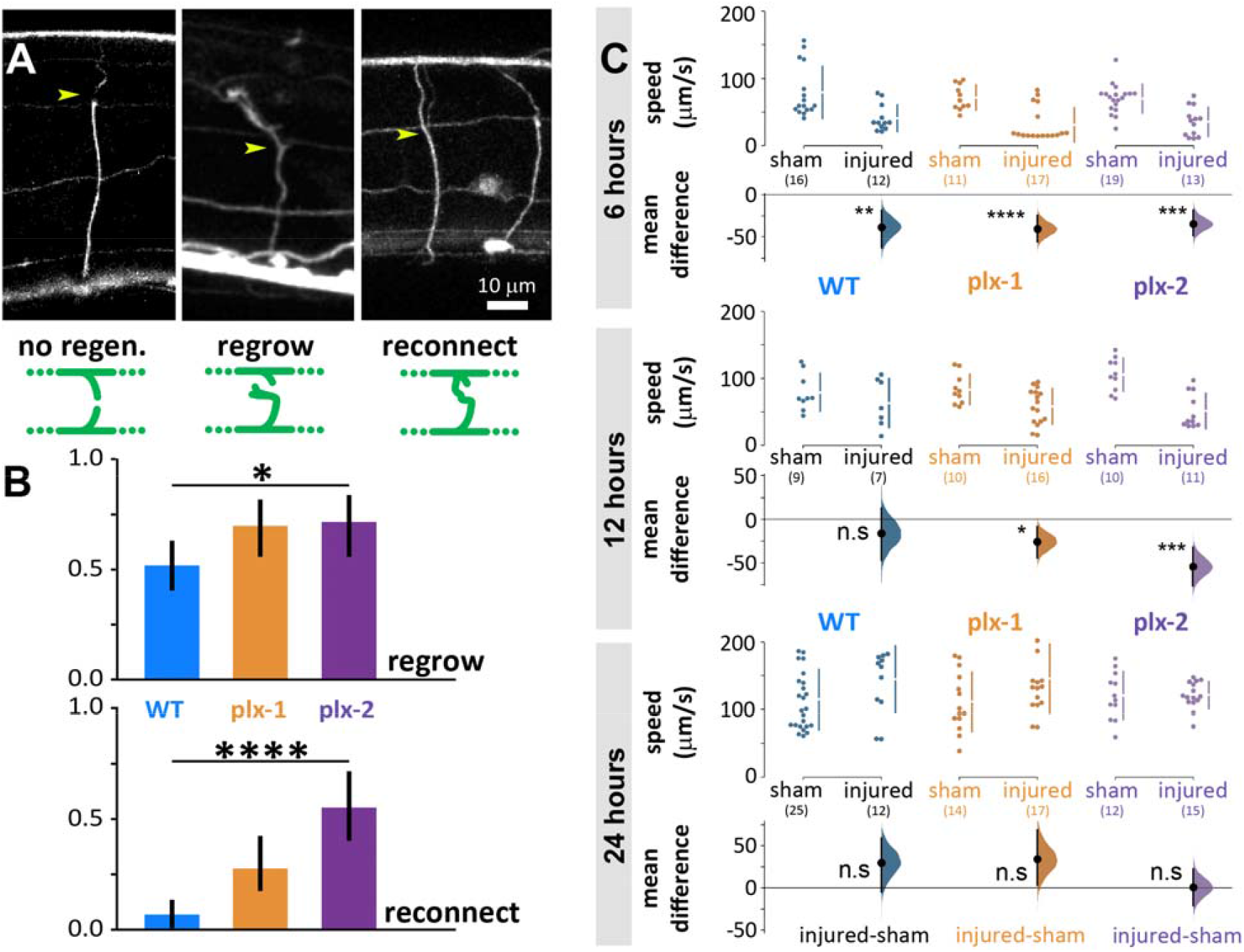
Neuronal regrowth and reconnection increased in the absence of plexins 24 hours after laser microsurgery, while locomotion speed fully recovers in all genotypes. A) We scored all commissural neurites 24 hours after microsurgery (yellow arrowhead for site of lesion, examples are 24 hours after lesion) and scored them as exhibiting either no-regeneration (WT), regrowth (*plx-2(ko)*, note growth cone), or reconnection (*plx-2(ko)*); schematically demonstrated in green diagrams, see methods. B) About half of wild type neurites regrew 24 h post-injury and only 7% reconnected. Both plexin knockout mutant strains exhibited more regrowth (top) and *plx-2* exhibited more reconnection (bottom, note that reconnection implies regrowth). Bars are fraction of observed neurites; ^*^p<0.05, ^****^p<0.0001; Fisher Exact on 2×3 contingency table. C) Injured animals of all groups moved significantly slower than sham operated 6 h post-injury, only wild type recovered at 12 h, and all genotypes recovered when compared to sham operated after 24 h. Data points are mean absolute translocation speed to both direction of locomotion; n.s. p>0.05, ^*^p<0.05 ^***^p<0.001,^****^p<0.0001; two-sided permutation t-test; in parentheses are the number of analyzed trajectories from 7-20 animals.

Six hours after microsurgery, wild type and mutant animals moved slower than sham-treated animals of the same genotype (sham vs injured: WT 79±39 vs 41±20 μm/s, p=0.004; *plx-1(ko)* 72±18 vs 31±25 μm/s, p<0.0001; *plx-2(ko)* 70±22 vs 35±21 μm/s, p=0.0001; Fig. 4C, top). Twelve hours after microsurgery, the mean locomotion speed of wild type animals has recovered to levels comparable to sham-treated, while mutant animals moved slower than their sham-treated controls (sham vs injured: WT 79±28 vs 63±36 μm/s; *plx-1(ko)* 84±22 vs 58±26 μm/s, p=0178; *plx-2(ko)* 106±25 vs 51±26 μm/s, p=0.0001; Fig. 4C, middle). Subsequently, 24 hours after microsurgery, mean locomotion speed has recovered to levels comparable to sham-treated animals for all groups (sham vs injured: WT 115±45 vs 145±49 μm/s; *plx-1(ko)* 111±44 vs 146±51 μm/s; *plx-2(ko)* 120±35 vs 121±20 μm/s; Fig. 4C, bottom).

## Discussion

Here we have demonstrated that the two plexins that mediate semaphorin signaling in *C. elegans* restrict neuronal regrowth and reconnection after injury. In their absence, injured neurons of plexin knockout mutants exhibit higher levels of regrowth (for both plexins) and reconnection (for *plx-2*).

By the nature of their ligands, the two plexins mediate different spatial signals. Paracrine interaction, such as those mediated by PLX-1 typically act at short-ranged by cell-to-cell interactions and conform subcellular resolution spatial information (Dalpé et al., 2004, 2005; Gurrapu and Tamagnone, 2016). Because both ligand and receptor are transmembrane proteins, the flow of information could be bidirectional, such as in the case of reverse-signaling through semaphorins, in which plexins function as ligands (Yu et al., 2010; Battistini and Tamagnone, 2016; Suzuki et al., 2022). On the other hand, juxtacrine interactions, such as those mediated by PLX-2 are spatially more disperse over tissue level where the ligand typically diffuses to set meaningful concentration gradients (Chen et al., 2007).

We demonstrated that neither the plexins nor the three semaphorins are necessary for gross neuromorphogenesis. However, at low penetrance their omission causes defasciculating and axon misguidance (Roy et al., 2000). In the nervous system, PLX-1 is only expressed by two motoneurons, namely DA8 and DA9, where it is involved in synaptic tiling during development by restricting the synaptic regions (Mizumoto and Shen, 2013). Because, to the most part, PLX-1 is expressed in muscle and other non-neuronal tissue (Fujii et al., 2002), we hypothesize that its restrictive effect on regeneration is achieved by interaction with the semaphorin SMP-1 presented by the motoneurons (Liu et al., 2005). The neurons could respond indirectly to the surrounding tissue via another signaling pathway, such as the ephrin pathway (as described for *efn-4* in relation to *plx-2/mab-20*; Nakao et al., 2007), or SMP-1 could mediate a direct cellular response via reverse-signaling from plexins to semaphorins (Yu et al., 2010; Battistini and Tamagnone, 2016; Suzuki et al., 2022). The other membrane-bound semaphorin, SMP-2, might not be involved in motoneuronal regeneration because it is not expressed by VNC motoneurons, but in body wall muscle and some sensory neurons in the head (Ginzburg et al., 2002). PLX-2 is expressed by four classes of motoneurons, and the most parsimonious hypothesis is that MAB-20 signals via PLX-2 to prevent aberrant neuronal regeneration; MAB-20 secretion from muscle cells generate a gradient that suppresses overgrowth of neurites in health and injury. A similar system was described for regenerating axons of murine spinal cord and brain, where expression of the receptor complex mediating SEMA3A function increases after injury, while SEMA3A secretion at the site of injury declines to undetectable levels during the period of axon regrowth, but persists to be secreted by cells adjacent to the injury site, creating an exclusion zone which regrowing axons do not penetrate (Pasterkamp and Verhaagen, 2001; Pasterkamp et al., 2001; de Winter et al., 2002). Notably, the absence of MAB-20 and PLX-2 had different effects on swimming speed, reminiscent of the different epidermal development phenotypes described for *mab-20(ko)* and *plx-2(ko)* (Nakao et al., 2007).

The phenotypes we describe for uninjured plexin and semaphorin knockout mutant animals are changes in speed and frequency of locomotion on agar surface and in liquid. To the most part, these effects are small in magnitude and include both increases and decreases compared to wild type animals. The largest effects were on the translocation speed of *smp-1(ko)* during swimming and even worse during crawling.

Because the semaphorin signaling pathways are involved in several aspects of embryonic development and its components are expressed in neuronal and non-neuronal tissue in the embryo, the phenotypes are likely the product of an accumulation of effects on structure and function of different tissue, such as muscle, cuticle, or the nervous system. Furthermore, the semaphorin pathways could regulate expression of downstream genes (Alto and Terman, 2017) that in turn effect locomotion behavior. Parsimoniously, because these effects are not the focus of this study, we removed the effect of these locomotion phenotypes by comparing animals after laser microsurgery to sham-operated animals of the same genotype. Moreover, the laser microsurgery experiments included only plexin knockout mutants and *smp-1(ko)* animals were not included in that comparison.

Locomotion behavior was impaired 6 hours post-injury and recovered back to pre-injury parameters 24 hours post-injury in wild type animals and both plexin knockout mutant animals. Because less than half of the neurites in the wild type animals regrew and only 0.07 reconnected, we hypothesize that the recovery is due to reorganization of the locomotion circuit to produce a meaningful motor pattern that is indistinguishable from that of an uninjured animal (Haspel et al., 2021). Similarly, the recovery of plexin knockout mutants that exhibit much higher levels of regrowth (for both) and reconnection (for *plx-2*) can be due to reorganization. Full recovery of locomotion with only partial recovery of neurites and synapses has been described in other systems (Oliphint et al., 2010), but the underlying circuit mechanism is unknown.

The conserved but concise semaphorin-plexin system and readily available genetic and transgenic tools in *C. elegans*, together with accurate injury and quick neuroregeneration and recovery of behavior provide an attractive experimental model. The secreted and membrane-bound semaphorin signaling pathways both restrict regeneration but in distinct processes that likely include spatial specificity and recurrent signals. Further studies, including of the effect on regeneration of each and combinations of the semaphorins and their localization, before and right after injury, as well as the spatiotemporal dynamics of related secondary messengers such as calcium and cAMP, will address proximate hypotheses about the involvement of semaphorin signaling in neural recovery from injury.

## Acknowledgements

We thank Dr Joseph Culotti (University of Toronto, Mt Sinai Hospital) and Dr Ricard Ikegami (UC Berkeley) for sharing the *plx-2* and *plx-1* reporter strains. We also thank Dr Kang Shen (Stanford University) and Dr Kota Mizumoto (University of British Columbia) for the *plx-1* reporter constructs. We thank Dr Monica Driscoll (Rutgers University) for her help in generating transgenic lines and the Hobert Lab (Columbia University) for the NeuroPal strains. We thank Joseph Soubany for help with locomotion anaysis. Research reported in this publication was supported by NINDS of the National Institutes of Health (1R15NS125565-01; MBH, SE, SB, TST, GH), by the State of New Jersey Commission on Spinal Cord Research (CSCR14ERG002; GH; and CSCR16IRG013; TST), and by the NJIT Undergraduate Research Initiative (URI; ES). Some strains were provided by the CGC, which is funded by NIH Office of Research Infrastructure Programs (P40 OD010440).

## Notes

### Competing Interest Statement

The authors have declared no competing interest.

### Summary of Updates

minor changes to the text replaced parts in all figures

